# Volumetric two-photon fluorescence imaging of live neurons using a multimode optical fiber

**DOI:** 10.1101/2020.04.27.063388

**Authors:** Raphaël Turcotte, Carla C. Schmidt, Martin J. Booth, Nigel J. Emptage

## Abstract

Multimode optical fibers (MMFs), combined with wavefront control methods, have achieved minimally-invasive *in vivo* imaging of neurons in deep-brain regions with diffraction-limited spatial resolution. Here, we report a method for volumetric two-photon fluorescence imaging with a MMF-based system requiring a single transmission matrix measurement. Central to this method is the use of a laser source able to generate both continuous wave light and femtosecond pulses. The chromatic spreading of pulses generated an axially elongated excitation focus, which we used to demonstrate volumetric imaging of neurons and their dendrites in live rat brain slices through a 60 μm-core MMF.

---

The ability to optically visualize neurons and their subcellular components across brain regions is integral to our understanding of neuronal function [1]. Volumetric two-photon fluorescence imaging methods based on axially extended focus are particularly well-suited to monitor neuronal dynamics occurring on the millisecond timescale [1–3]. By using a Bessel-like beam, a volume can indeed be imaged within a single frame through integration of the signal across depth while maintaining a lateral resolution sufficient for resolving synaptic components, such as dendritic spines [3]. Volumetric imaging approaches have been developed for micro-endoscopy with multicore optical fiber [4] and used to monitor calcium transients in deep-brain regions (e.g. hypothalamus) with gradient index (GRIN) lenses [5]. Bessel-like beams have also been used for one-photon fluorescence in light-sheet microscopy with MMFs [6].

MMFs have emerged as a central component of a minimally-invasive micro-endoscope technology owing to their small size relative to other devices [7]. Indeed, their diameter, between 60-150 μm, enables direct insertion several millimeters into the living mammalian brain while causing minimal tissue damage and, hence, is likely to minimally affect the neuronal network [7–9]. An essential component of MMF technology is wavefront control, required to circumvent the random phase delays and mode coupling that occur inside a fiber [10, 11]. By shaping the wavefront entering a MMF using a spatial light modulator (SLM), a focused point of light can be formed at the distal end of the fiber, suitable for the excitation of fluorophores in a sample. Three-dimensional fluorescence imaging through MMF requires spatiotemporal focusing of femtosecond pulses [12]. The spectral content of these pulses must also be carefully considered as MMF-induced perturbations are wavelength dependent [13, 14]. With these approaches, multiphoton imaging through MMFs has been demonstrated with beads samples, CHO cells *in vitro*, and fixed cochlear hair cell samples [15–18].

In this Letter, we demonstrate, for the first time to our knowledge, two-photon fluorescence imaging with an axially extended focus through a MMF. We employed a step-index MMF offering a relatively high numerical aperture of 0.37 and a total diameter of 75 μm, which are optimal in terms of maximizing lateral resolution and minimizing invasiveness, respectively. In addition, we took advantage of the wavelength-dependent axial shift caused by the limited bandwidth of step-index MMFs [14] to generate an extended focus through chromatic spreading, and showed that this focus can be used for volumetric imaging of neurons in living rat brain slices.

A schematic of the experimental system is shown in Fig. 1. The light from a dual-source laser (Mira Optima 900-F V5 XW, Coherent, 830 nm, 76 MHz, mode: continuous wave (CW) or femtosecond pulses (pulsed, 146.5 ± 0.5 fs)) was separated into two paths within the source module using a polarizing beam splitter (PBS). In the first path, the light was directed to the imaging module using free-space optics. In the second path, the light was coupled into a single mode fiber (SMF) and brought to the calibration module. Light entering the imaging module was delivered to a liquid-crystal spatial light modulator (LC-SLM, Meadowlark Optics, HSPDM 512, 512 × 512 pixels). A half-wave plate (HWP) was located before the telescope in the source module to maximize the power into the first-order diffraction beam. The first-order diffraction beam from the LC-SLM carried the shaped wavefront and was aligned on-axis such that an iris positioned at the conjugated plane (L1, focal length (FL): 100 mm) filtered out all other diffraction orders. A dichroic mirror (DM, Semrock, FF750-SDi02-25×36) positioned immediately after the iris reflected the illumination light toward another lens (L2, FL: 50 mm) that created an intermediate SLM plane and a quarter-wave plate (QWP) made the polarization circular. The light from this intermediate plane was coupled (L3, FL: 8 mm) into a MMF. We used a step-index MMF having a numerical aperture of 0.37, a core diameter of 60 μm and a total diameter of 75 μm (Doric Lenses, MFC_060/075-0.37_20mm_ZF1.25_FLT). The total diameter of this implant is ideal for minimally invasive *in vivo* applications compared to other implants such as GRIN lenses usually in the range of 0.5 - 1 mm [7, 8].

**Fig. 1.**
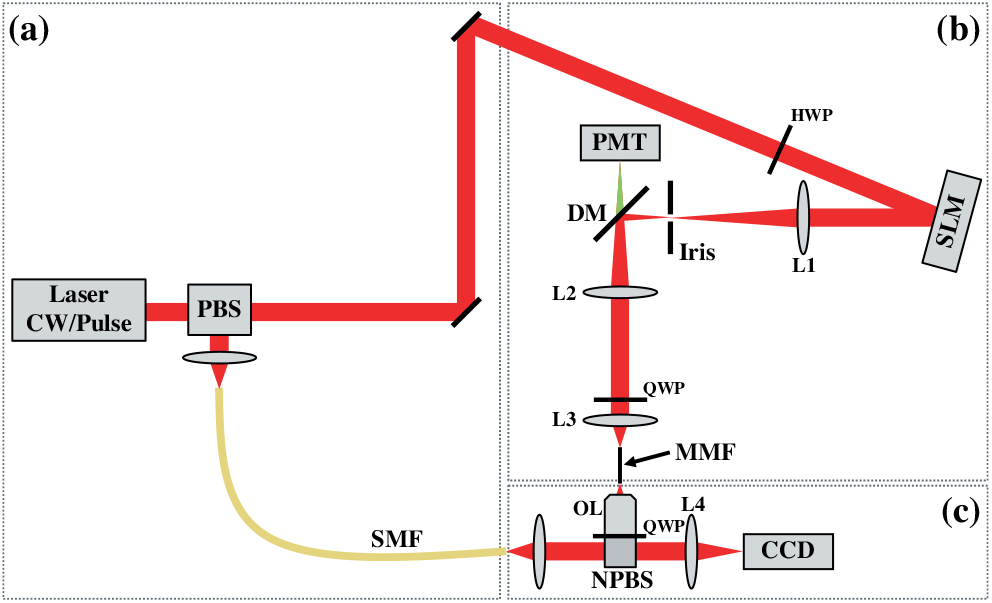
Schematic of the experimental system. (a) The source module contains a laser able to generate CW and pulsed outputs. (b) The imaging module includes an LC-SLM for wave-front control during imaging and calibration, as well as a PMT for detection of the epi-propagating fluorescence. (c) The calibration module is used for recording illumination focus data and for evaluating the transmission matrix.

The removable calibration module was used in the evaluation of the transmission matrix (TM), which describes the complex field transformation through the MMF, by imaging a selected plane some distance away from the MMF distal facet while it interfered with the SMF reference beam [19]. This selected calibration plane was relayed by a microscope objective lens (OL, Olympus, RMS20X, NA = 0.4) and an achromatic doublet (L4, FL: 150 mm) onto a charge-couple device camera (CCD, Basler pilot, piA640-210gm). In between the lenses, the signal was converted back into the linear polarization state using a QWP and merged with a reference signal using a 50:50 non-polarizing beam-splitter (NPBS). The TM was evaluated with the laser source in CW mode as described in details elsewhere [7, 20]. For all data, the calibration was performed 40 μm from the distal facet. Two-photon fluorescence images were formed by digital raster-scanning with the LC-SLM [7, 20], i.e. by presenting sequentially a distinct wavefront for each pixel in the final image with the laser in pulsed mode. The wavefront was shaped using information from the TM to focus light and excite fluorophores. The emitted fluorescence was collected by the MMF, collimated by L3, and focused by L2, through the dichroic mirror and other spectral filters (Semrock, FF01-750/SP25, FF01-607/70-25, and BLP02-561R-25), onto a photomultiplier tube (PMT).

The performance of our system was first assessed by generating foci at the MMF distal end for multiphoton fluorescence excitation. For this purpose, illumination foci were sequentially captured with the CCD camera in the calibration module for both laser operating modes (Fig. 2). The TM was used to calculate the LC-SLM pattern for each focus in an equidistant output grid pattern of 13 × 13. Only half of the foci are shown to ensure that there is no overlap between adjacent foci. The recording of focus data was accelerated by using the graphical processing unit for grabbing images recorded with the CCD camera [20]. Width measurements were made using a custom MATLAB script and employing a numerical approach to evaluate the full width at half maximum and 1/e^2^ width.

**Fig. 2.**
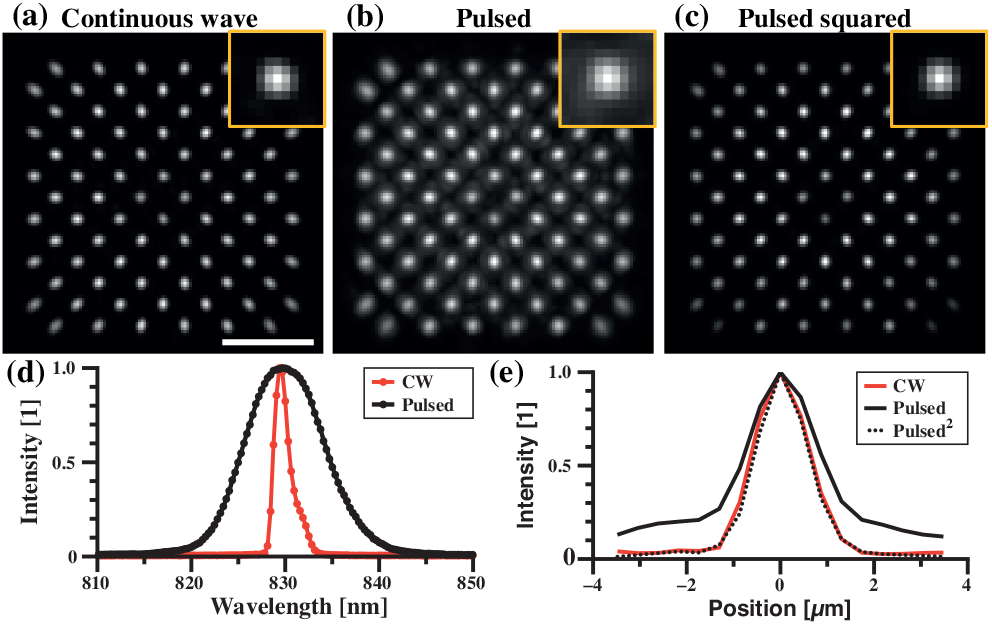
The non-linearity of two-photon fluorescence improves focusing quality by suppressing background. (a,b) Illumination foci achieved with wavefront control through a MMF and recorded sequentially at different lateral positions (composite maximum intensity projection from images of individual foci) when the laser is operating in (a) CW mode and (b) pulsed mode. (c) Squared of the intensity distribution shown in (b) to illustrate the two-photon illumination. Scale bar: 15 μm; inset width: 4.8 μm. (d) Spectra of the laser source in CW and pulsed mode. (e) Intensity profile for the inset foci shown in (a-c).

With CW light, a focal light distribution was achieved in the center of the fiber (full width at half maximum (FWHM): 1.4 ± 0.1 μm; 1/e^2^ width: 2.3 ± 0.2 μm). A maximum intensity projection of illumination foci recorded at different lateral locations revealed that the performance was maintained across the field-of-view (Fig. 2(a)), that is, uniform illumination was achieved. With the calibration performed in CW mode, it was then necessary to switch to pulsed mode to generate multiphoton fluorescence. The change in the laser temporal mode (308 ± 13 fs) had minimal impact on the lateral illumination foci despite the high sensitivity of the transmission matrix to perturbations and spectral changes (Fig. 2(b)) [13, 14]. With the laser in pulse mode the width of the illumination foci increased (FWHM: 1.7 ± 0.1 μm) and tapered off more slowly at the edges than for the CW mode (Fig. 2(e); 1/e^2^ width: 3.4 ± 0.3 μm). Some deterioration in the focus quality was expected because calibration is wavelength dependent and spectral broadening would occur on transitioning from CW to pulsed mode (Fig. 2(d); FWHM CW: 1.9 ± 0.3 nm; FWHM pulse: 9.8 ± 0.3 nm). However, it is important to recognize that as the intensity of two-photon fluorescence is dependent on the square of the photon distribution, then the square of the illumination foci should be evaluated to characterize the two-photon fluorescence excitation profile (Fig. 2(c)). When assessed in this way, a similar point quality is achieved in pulsed mode as for the CW mode, with a FWHM of 1.3 ± 0.1 μm with almost entirely suppressed extended edges (Fig. 2(e); 1/e^2^ width: 2.3 ± 0.2 μm). This non-linear background reduction is analogous to side ring suppression in two- and three-photon Bessel foci [21]. Thus, we have successfully implemented transverse spatial focusing of femtosecond laser pulses through a MMF.

We then assessed the axial characteristic of the foci by repeating the above illumination measurements at different distal planes by translating the calibration module in increments of 2 μm (Fig. 3). The axial FWHM with CW light was limited to 13.6 ± 1.1 μm (Fig. 3(a,d)). With pulsed light, the axial FWHM increased by a factor of 2.2× to 29.6 ± 0.7 μm (Fig. 3(b,d)). As expected the increased spectral content of pulses caused an elongation of the light distribution [14]. The axial FWHM for the two-photon fluorescence excitation volume was computed to be 24.4 ± 0.6 μm (Fig. 3(c,d)). The lateral FWHM as a function of the distal location revealed that in CW mode the lateral focus was preserved over a relatively short distance, as expected for Gaussian focusing, and that this distance was substantially extended upon switching the laser in pulsed mode (Fig. 3(e)). Importantly, the lateral FWHM was maintained below 1.5 μm over a distance of 40 μm for the pulsed squared case which is sufficient for volumetric imaging (Fig. 3(e)).

**Fig. 3.**
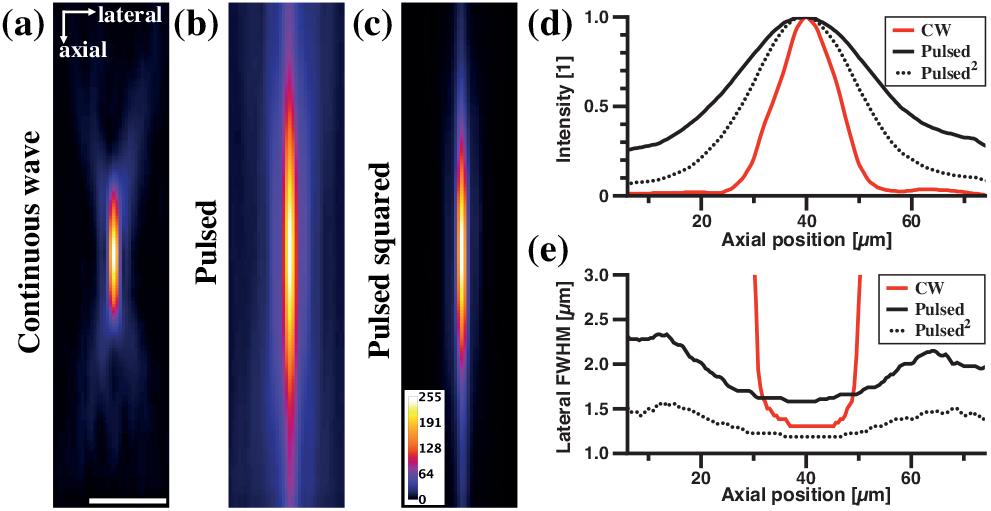
Spectral dispersion axially elongates the focus when switching the laser from CW to pulsed mode. (a,b) Illumination foci achieved with wavefront control through a MMF and recorded sequentially at different axial positions (axial-lateral profile) when the laser is operating in (a) CW mode and (b) pulsed mode. (c) Square of the intensity distribution shown in (b) to illustrate the two-photon excitation. (d) Intensity as a function of the axial position (distance from the distal facet) for the data shown in (a-c). (e) Lateral FWHM as a function of the axial position for the data shown in (a-c). Scale bar: 10 μm.

Having established a procedure for generating the desired light distribution in the sample, we tested the two-photon fluorescence imaging capability of the MMF system using fluorescent beads. Figure 4(a) shows an image of 2-μm red fluorescent beads (Invitrogen, FluoSphere, red 580/605) acquired at a distance of 40 μm from the distal facet. An excellent signal-to-noise ratio was achieved (SNR = 6.0 ± 0.6) with a total incident power at the sample of 10 mW. The fluorescence background from the focal plane has been a limiting factor for attaining high dynamic range imaging with linear fluorescence. This is true although the peak intensity can easily reach values 300 time larger than the average background illumination [9, 22]. Indeed, the background fluorescence from bright out-of-focus objects can easily overwhelm the signal from a dim in-focus object considering that the fluorescence background is integrated over a relatively large volume. Further suppression of the in-focus background from the non-linearity of two-photon fluorescence would therefore be beneficial for increasing the dynamic range of the imaging system. In our system, we obtained an enhancement factor, evaluated as the ratio of the peak to average intensity, of 383 ± 34 in CW mode and 63 ± 6 in pulsed mode. The calculated enhancement factor for the pulsed squared was of 914 ± 114. The large enhancement factor for pulsed squared is consistent with the minimal background present while performing two-photon fluorescence imaging. Figure 4(b) shows an axial-lateral profile of single beads recorded by translating the sample in steps of 2 μm. This profile reveals the extended focus along the axial dimension as signal was detectable even when the sample was located 30 μm away from the calibration plane. The lateral FWHM of a single bead was relatively constant (2.2 ± 0.1 μm) in the 30 μm around the calibration plane (Fig. 4(c)).

**Fig. 4.**
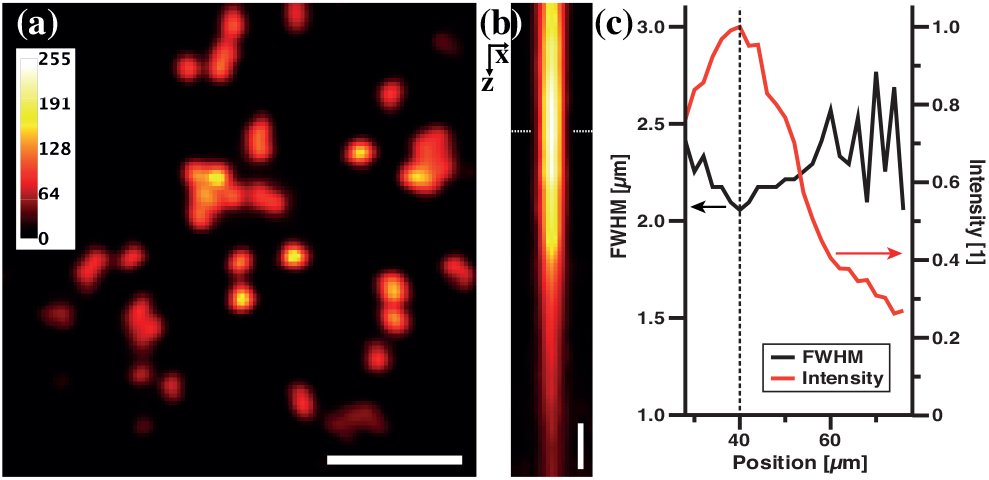
MMF volumetric two-photon fluorescence imaging of beads (2 μm diameter, red 580/605). (a) Image of beads located at the calibration plane (40 μm from the MMF facet). Scale bar: 15 μm. (b) Averaged axial(z)-lateral(x) profile from 5 beads. Images were acquired by translating the sample in steps of 2 μm. Scale bar: 5 μm. (c) Intensity profile and FWHM as a function of the distance between the distal facet and sample for the data shown in (b). Dash lines indicate the calibration plane.

As our design is optimized for imaging in live-tissue we assessed its performance by imaging Alexa Fluor 594 labelled neurons in living brain slices from the rat hippocampus. Organotypic hippocampal slices were prepared from male Wistar rats (P7-P8; Harlan UK) and imaged in physiological Tyrode’s solution (in mM: 120 NaCl, 2.5 KCl, 30 glucose, 2 CaCl_2_, 1 MgCl_2_, and 25 HEPES; pH = 7.2-7.4) at room temperature. Fluorescent structures were readily located and could be identified as neuronal cell soma (Fig. 5(a)). Pyramidal cell apical dendrites were also clearly visible projecting from the soma. In an attempt to visualize smaller features, we followed the apical dendrite away from the cell body and were able to detect secondary dendritic branching from the primary apical dendrite (Fig. 5(b)). Visualizing live cells within tissue is an important application of multiphoton MMF imaging [16–18] and our optical system was optimized for live animal imaging through compactness, stability, and mobility [7, 20]. Indeed, the modular design of the system is particularly well suited for *in vivo* applications and we expect that this method will enable volumetric imaging of neurons in living mammalian brains once implemented at optimized wavelengths.

**Fig. 5.**
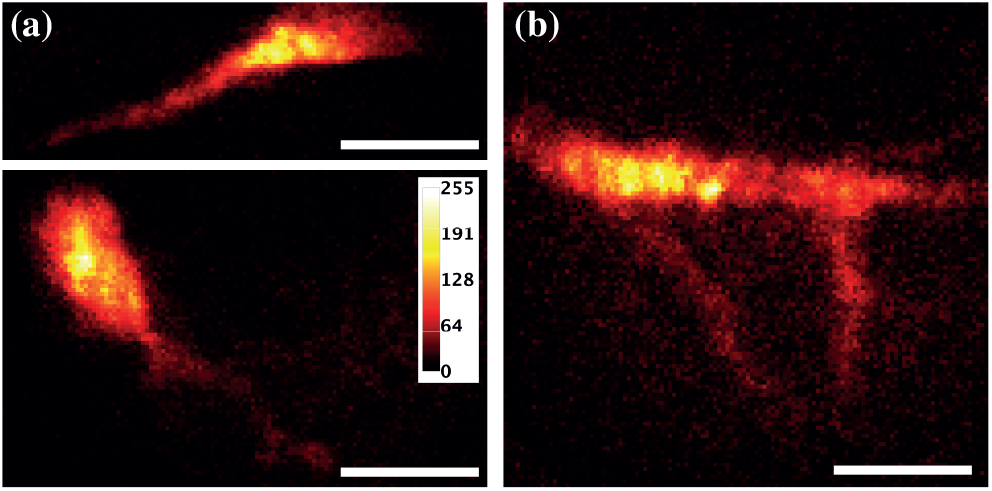
MMF volumetric two-photon fluorescence imaging in live brain tissue from the rat hippocampus reveals. (a) neuronal cell bodies and apical dendrites as well as (b) smaller dendrites branching off from an apical dendrite, after the labeling CA3 neurons with 5 mM of Alexa Fluor 594 using whole-cell patch electrophysiology. Scale bar: 15 μm.

In summary, we have achieved volumetric multiphoton imaging of neuronal soma and associated dendrites in living brain tissue using wavefront control through a MMF. This demonstration was accomplished by integrating a dual CW/pulsed laser source into a micro-endoscopic system. A single calibration with the laser in CW mode was sufficient to determine the TM needed to generate an chromatic extended depth of field for femtosecond pulses. Indeed, dispersion in the step-index MMF axially distributed the broader spectral content of pulses compared to the CW light around the calibration plane, effectively generating an axially elongated chromatic focus [23, 24]. The use of a CW laser for the calibration had the added benefit of allowing a design employing removable distal optics, an important advantage for biological imaging applications. Finally, while the non-linearity of the multiphoton process decreased the background, it also exacerbated spatial non-uniformities in the illumination. However, these non-uniformities are independent of the calibration method as they arise intrinsically from the fiber geometry and could be alleviated by performing spatially-variant deconvolution in order to regularize the spatial response [25].

## FUNDING

Biotechnology and Biological Sciences Research Council (BBSRC) (BB/P0273OX/1); European Research Council Advanced Grant AdOMIS (695140).

## Disclosures

The authors declare no conflicts of interest.

## Notes

### Competing Interest Statement

The authors have declared no competing interest.

### Summary of Updates

Additional data was included related to the characterisation of the axial dimension and the manuscript re-written accordingly.

## REFERENCES

1. N. Ji, J. Freeman, and S. L. Smith, Nat. Neurosci. 19, 1154 (2016).

2. G. Thériault, M. Cottet, A. Castonguay, N. McCarthy, and Y. D. Koninck, Front. Cell. Neurosci. 8, 139 (2014).

3. R. Lu, W. Sun, Y. Liang, A. Kerlin, J. Bierfeld, J. D. Seelig, D. E. Wilson, B. Scholl, B. Mohar, M. Tanimoto, M. Koyama, D. Fitzpatrick, M. B. Orger, and N. Ji, Nat. Neurosci. 20, 620 (2017).

4. A. Orth, M. Ploschner, I. S. Maksymov, and B. C. Gibson, Opt. Express 26, 6407 (2018).

5. G. Meng, Y. Liang, S. Sarsfield, W.-c. Jiang, R. Lu, J. T. Dudman, Y. Aponte, and N. Ji, eLife 8, 4546 (2019).

6. M. Plöschner, V. Kollárová, Z. Dostál, J. Nylk, T. Barton-Owen, D. E. K. Ferrier, R. Chmelík, K. Dholakia, and T. Čižmár, Sci. Reports 5, 435 (2015).

7. S. A. Vasquez-Lopez, R. Turcotte, V. Koren, M. Plöschner, Z. Padamsey, M. J. Booth, T. Čižmár, and N. J. Emptage, Light. Sci. & Appl. 7, 110 (2018).

8. S. Turtaev, I. T. Leite, T. Altwegg-Boussac, J. M. P. Pakan, N. L. Rochefort, and T. Čižmár, Light. Sci. & Appl. 7, 92 (2018).

9. S. Ohayon, A. Caravaca-Aguirre, R. Piestun, and J. J. DiCarlo, Biomed. Opt. Express 9, 1492 (2018).

10. T. Čižmár and K. Dholakia, Opt. Express 19, 18871 (2011).

11. R. Leonardo and S. Bianchi, Opt. express 19, 247 (2011).

12. E. E. Morales-Delgado, D. Psaltis, and C. Moser, Opt. Express 23, 32158 (2015).

13. M. Mounaix, D. M. Ta, and S. Gigan, Opt. Lett. 43, 2831 (2018).

14. T. Pikálek, J. Trägårdh, S. Simpson, and T. Čižmár, Opt. Express 27, 28239 (2019).

15. E. E. Morales-Delgado, S. Farahi, I. N. Papadopoulos, D. Psaltis, and C. Moser, Opt. Express 23, 9109 (2015).

16. S. Sivankutty, E. R. Andresen, R. Cossart, G. Bouwmans, S. Monneret, and H. Rigneault, Opt. Express 24, 825 (2016).

17. E. Kakkava, M. Romito, D. B. Conkey, D. Loterie, K. M. Stankovic, C. Moser, and D. Psaltis, Biomed. Opt. Express 10, 423 (2019).

18. J. Trägårdh, T. Pikálek, M. Šerý, T. Meyer, J. Popp, and T. Čižmár, Opt. Express 27, 30055 (2019).

19. S. M. Popoff, G. Lerosey, R. Carminati, M. Fink, A. C. Boccara, and S. Gigan, Phys. Rev. Lett. 104, 100601 (2010).

20. M. Plöschner and T. Čižmár, Opt. Lett. 40, 197 (2015).

21. C. Rodríguez, Y. Liang, R. Lu, and N. Ji, Opt. Lett. 43, 1914 (2018).

22. R. Turcotte, C. C. Schmidt, N. J. Emptage, and M. J. Booth, Opt. Lett. 44, 2386 (2019).

23. O. Cossairt and S. Nayar, “Spectral Focal Sweep: Extended depth of field from chromatic aberrations,” in 2010 IEEE ICCP, (Cambridge, MA, USA), pp. 1–8.

24. N. M. Fitzgerald, C. Dainty, and A. V. Goncharov, Opt. Express 25, 31696 (2017).

25. R. Turcotte, E. Sutu, C. C. Schmidt, N. J. Emptage, and M. J. Booth, Biomed. Opt. Express 11, 4759 (2020).

